# Exploration of Plant Biodiversity to Investigate Regulation of Photosynthesis in Natural Environment

**DOI:** 10.1101/368852

**Authors:** Simone Sello, Alessandro Alboresi, Barbara Baldan, Tomas Morosinotto

## Abstract

Photosynthesis is regulated in response to dynamic environmental conditions to satisfy plant metabolic demand but also to avoid over-excitation of the electron transport chain generating harmful reactive oxygen species. Photosynthetic organisms evolved several mechanisms to modulate light harvesting and electron transport efficiency to respond to conditions changing at different timescales, going from fast sun flecks to slow seasonal variations.

These regulatory mechanisms changed during evolution of photosynthetic organisms, also adapting to various ecological niches. The investigation of plant biodiversity is valuable to uncover conserved traits and plasticity of photosynthetic regulation. In this work a set of plants belonging to different genera of angiosperms, gymnosperms, ferns and lycophytes was investigated by monitoring their photosynthetic parameters in different seasons, looking for common trends and differences. In all plants analysed photosynthetic electron transport rate was found to be modulated by growth light intensity, ensuring a balance between available energy and photochemical capacity. Growth light also influenced the threshold where heat dissipation of excitation energy, also called Non-Photochemical Quenching (NPQ), was activated. On the contrary NPQ amplitude did not correlate with light intensity experienced by the plants but was a species-specific feature.

NPQ zeaxanthin-dependent component, qZ, was found to be the most variable between different plants, modulating the intensity of the response but also the kinetic properties of its activation and relaxation. The slower NPQ component, qI, was instead found to be uncorrelated with photoinhibition eventually suffered by plants.

## INTRODUCTION

Photosynthetic organisms exploit sunlight to power the transfer of electrons from water to NADP^+^ thanks to the activity of two photosystems (PS), PSI and PSII. Electron transport yields into the synthesis of ATP and NADPH that are exploited to drive CO_2_ fixation. Photosynthesis is a complex metabolic pathway requiring a fine regulation to respond to natural environment with highly variable conditions. In limiting light conditions, available energy should in fact be exploited with maximal efficiency to support metabolism demand. On the contrary, when incident radiation exceeds plant electron transport capacities the activation of protection mechanisms is required to minimize reactive oxygen species (ROS) production and photoinhibition (Eberhard *et al*., 2008; Li *et al*., 2009). Mechanisms for regulation of photosynthesis needs to respond not only to illumination but also to other abiotic parameters like temperature, water and nutrient supply that directly affect photosynthetic reactions and/or ATP/NADPH demand (Külheim *et al*., 2002; Peltier *et al*., 2010; Allahverdiyeva *et al*., 2014).

The regulation of photosynthesis under variable conditions influences plant fitness in natural environments (Külheim *et al*., 2002) as well as crops productivity in cultivated fields (Kromdijk *et al*., 2016). Changes in light intensity are a very common occurrence in outdoor environment with different timescales spanning from seconds, when clouds or the overlaying canopy interfere with the incident sunlight, up to weeks, following seasonal changes. For this reason plants evolved multiple regulatory mechanisms, with different activation timescales, enabling responses to the short- and long-term variations in environmental cues (Walters, 2005; Eberhard *et al*., 2008; Li *et al*., 2009). Within seconds upon an increase in light intensity, the thermal dissipation of excess absorbed energy is activated with a mechanism called Non Photochemical Quenching (NPQ). In vascular plants NPQ activation depends on the presence of the protein PSBS, activated by a decrease in lumenal pH (Li *et al*., 2000). In the timescale of minutes, strong illumination activates the synthesis of the xanthophyll zeaxanthin from violaxanthin by activation of the enzyme violaxanthin de-epoxidase (VDE) (Arnoux *et al*., 2009; Nilkens *et al*., 2010). Zeaxanthin, besides enhancing NPQ and retarding its relaxation in the dark, also plays a direct role in ROS scavenging (Havaux & Niyogi, 1999). On a longer-term, plants respond to changing light intensity by activating an acclimation response consisting in the modulation of gene expression, protein content and thylakoid membrane ultrastructure to acclimate to the changing environmental conditions (Walters, 2005; Eberhard *et al*., 2008). As example, the concentration of carbon fixing enzymes increases in high light acclimated plants, allowing for an increased exploitation of the available radiation.

Photosynthetic electron transport is also regulated in response to environmental changes and under strong illumination Cyt b_6_f activity was shown to be inhibited in a response called photosynthetic control that reduces electron transport flow (Joliot & Johnson, 2011). Electron transport is also regulated by activation of alternative electron transport pathways in addition to the main linear transport from water to NADP+ that protect PSI from over-reduction, also contributing to balancing ATP / NADPH production (Peltier *et al*., 2016; Yamori & Shikanai, 2016).

Most of the available knowledge on molecular mechanisms of photosynthesis regulation originates from studies with model organisms, mainly *Arabidopsis thaliana*. In recent years, however, several examples showed that these regulatory strategies may be diversified in different organisms. As example, study of NPQ activation mechanisms in the lycophyte *Selaginella martensii* suggested involvement in this organism of a specific phosphorylation of antenna proteins for regulation of both photosystems (Ferroni *et al*., 2017). Regulation of electron transport involving flavodiiron proteins has instead been shown to be present in all photosynthetic organisms but Angiosperms (Ilík *et al*., 2017). Several other emerging examples support the idea that exploring natural biodiversity is a valuable opportunity to obtain information on the mechanisms adopted by plants for the regulation of photosynthesis and to identify both common and specific features. In this work we investigated the mechanisms for regulation of light phase of photosynthesis in 20 different plant species, followed during different seasons of the year to assess their response to different growing conditions, evidencing common strategies for photosynthesis regulation.

## MATERIALS AND METHODS

### Selection of plant species and sampling

The choice of species to include in the analysis was based following the distribution of organisms in two recently published phylogenetic trees for gymnosperms and angiosperms (Figure S1) (Soltis *et al*., 2011; Lu *et al*., 2014). Among the species available at the local Botanical Garden (http://www.ortobotanicopd.it/en/) some non-flowering plants were chosen: *Cycas revoluta* and *Zamia fischeri* as Cycadaceae, *Ginkgo biloba* as Ginkgoaceae, *Taxus baccata* and *Cephalotaxus harringtonii* in the macrogroup formed by Taxaceae and Cephalotaxaceae, *Taxodium distichum, Sequoia sempervivens*, and *Tuja occidentalis* as Cupressaceae. Among Angiosperms, two monocotyledons, *Saccharum officinarum* and *Musa* sp., and six dicotyledons, *Piper nigrum* as Magnoliidae, *Coffea arabica* as Gentianales, *Camellia sinensis* as Ericales, *Salvia divinorum* as Lamiales, *Liquidambar orientalis* as Saxifragales and *Robinia pseudacacia* as Fabales, were chosen. Four non-seed plants were also included, two lycophytes belonging to the genus *Selaginella (S. kraussiana and S. pallescens)*, and two ferns, *Tectaria gemmifera* and *Diplazium esculentum* (Chaw *et al*., 2000). Samplings were carried out in different time points in 2016 and 2017 to assess also the effect of acclimation to different seasons and in total at least 10 independent measurements were performed per each species. Technical reproducibility was verified to validate our procedures to be small and therefore to have limited impact as cpompared with seasonal variance (Figure S2). Samples were collected and dark-adapted for 45 minutes at the sampling site to maintain growth temperature conditions. Sun-exposed leaves were chosen for measurements when possible, always avoiding young or old leaves. In the case of *C. revoluta, Z. fischeri, R. pseudacacia*, that have fronds or composed leaves, primary veins were avoided during photosynthetic measurements. In the case of needle-shaped leaves, tip needles were avoided. When microphylls or needle leaves were used, multiple samples were aligned to cover the measuring area of detector heads. In all cases, brownish parts, pathogen-attacked leaves, yellow spots and every other visually detectable deviation from healthy leaves were discarded.

Information about light intensity was collected at the time of the measurement using a LI-COR LI-250A light meter (Biosciences). Measurements repeated 3 times between 10.30 and 12.30. Temperature at sampling site was also monitored.

### Measurement of photosynthetic parameters

Chl fluorescence and PSI absorbance were monitored using a Dual-PAM 100, every analysis lasted 40 min in total. A measuring light was set to 5 μmol photons m^-2^ s^-1^, saturation pulse was 6000 μmol photons m^-2^ s^-1^ for 600 ms. After 45 min of dark adaptation to completely oxidize the photosynthetic electron transport chain, samples were treated with actinic light of increasing intensity (7, 14, 27, 70, 130, 175, 225, 276, 330, 415, 531, 653, 811, 1007, 1275, 1583, 2001 μmol photons m^-2^ s^-1^). After 60 seconds at one light intensity, a saturation pulse was applied to assess the redox state of PSII and PSI and then a more intense actinic light was applied for 60 seconds. When actinic light was switched off, leaves were left in the dark for 20 minutes, with regular application of saturation pulses. PSII and PSI parameters were calculated as follows: Fv/Fm as (Fm – Fo)/Fm, Y(II) as (Fm’ – F)/Fm’, NPQ as (Fm – Fm’)/Fm’, ETRII as Y(II) × PPFD × αII, Y(I) as 1 –Y(ND) – Y(NA), Y(NA) as (Pm – Pm’)/Pm, Y(ND) as (1 – P700 red), ETRI as Y(I) × PPFD × αI (Maxwell & Johnson, 2000).

### Data processing

Collected data were processed to obtain quantitative parameters describing the recorded kinetics. NPQ induction curves were fitted with a sigmoidal function (Hill function (y=V_max_*x_n_*(k_n_+x_n_)-1) found to be the most suitable to describe its activation in leaves treated with increasing actinic light intensities, as already observed in algae (Bernardi *et al*., 2016). Three main parameters were obtained, V_max_ that is the maximal NPQ value reached during the experiment, k is the light intensity needed to activate 2/3 of the NPQ; and n is a parameter describing the sigmoidicity of the curve. Induction kinetics of all other photosynthetic parameters considered (ETRI, ETRII, YI, YII, YND, YNA) were fitted using a similar approach, using a Hill function for induction, thus obtaining Vmax, k and n for each analysis.

Following earlier examples in the literature (Nilkens *et al*., 2010; Holzwarth *et al*., 2013), NPQ relaxation was instead fitted with a biexponential decay (y = y_0_ + A_1_ e-x/t_1_ + A_2_ e -x/t_2_). Y_0_, the minimum value reached by the fitting, represented the NPQ that is not relaxed after 20 minutes of dark recovery and thus represents an estimation of the slow qI component. A_1_ and A_2_ instead corresponded to the fastest qE and qZ components.

### Statistical analysis

The presence of correlations between all environmental and photosynthetic parameters collected was analysed with Pearson’s correlation (Yin *et al*., 2017) using SPSS software v.22 (Chicago, IL, USA). Bonferroni correction was considered to correct the *α*-value when several dependent or independent statistical tests were performed simultaneously. This correction gives the advantage to avoid spurious positives. For this method, the *α*-value (0.01) was divided by the number of variables (33), and the new value is used as the new α. Correlations with lower *p*-values than the new α were not considered for further analyses. Linear regression analyses were used to examine significant correlations of traits.

## RESULTS

### Measurement of photosynthetic parameters sampling biological and environmental variability

Plants from 20 different species were selected among Angiosperms, Gymnosperms, Lycophytes and ferns to represent available biodiversity (Figure 1). Photosynthetic parameters were measured in different seasons for approximately two years, to assess also the eventual influence of acclimation to seasonal conditions. In some cases, it was also possible to measure leaves either directly exposed to sunlight or in the canopy shade to increase variability of environmental conditions.

**Figure 1.**
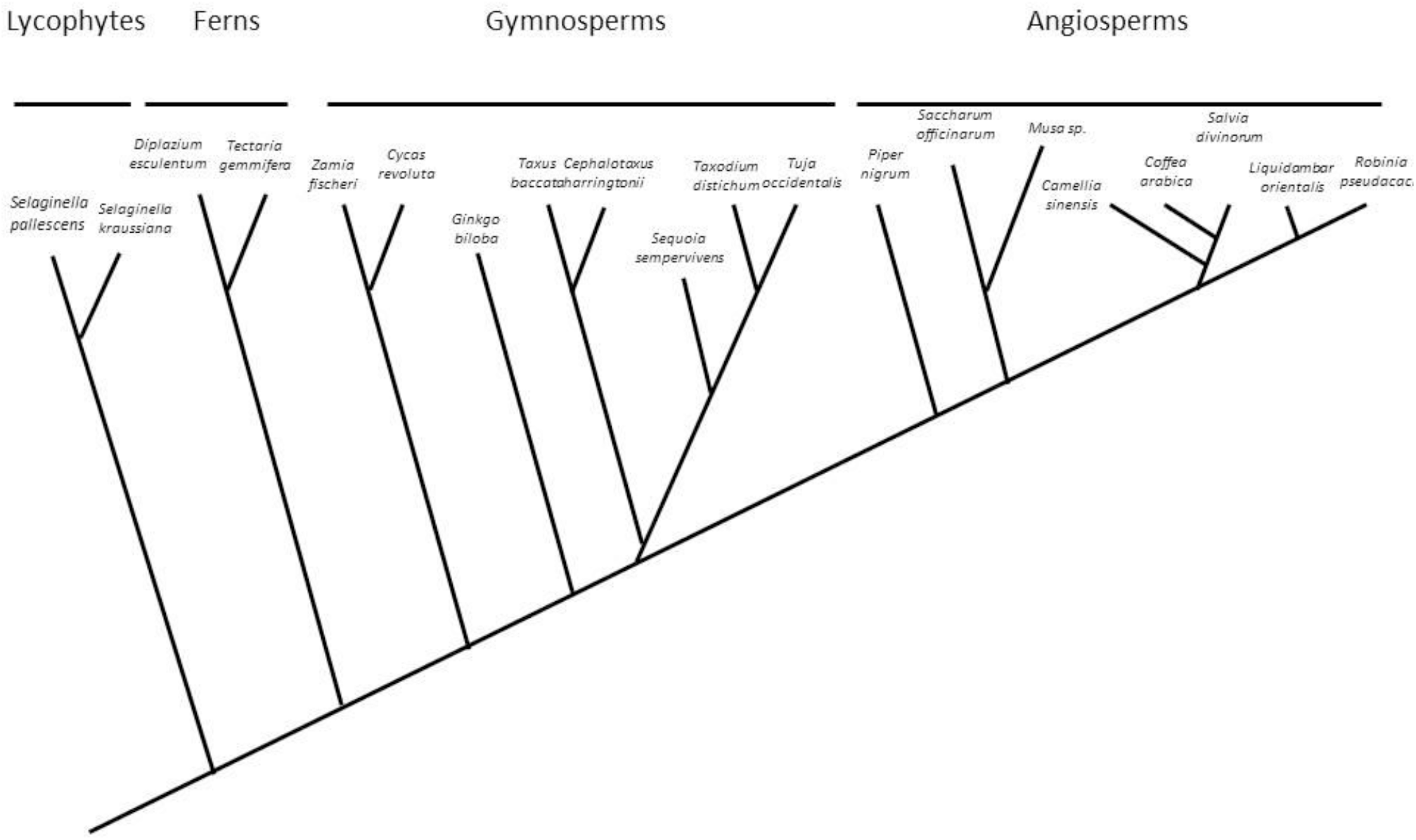
Plant species selected for the present work. Phylogenetic relationship between species selected for this work represented with a simplified tree, based on data from Vasco et al., Front. Plant Sci., 2013 for lycophytes and ferns; Ying Lu, et al, PLOS One 2014 for gymnosperms; Soltis et al., American J Bot, 2011 for angiosperms. The lengths of the branches are not indicative of time separation

Plant ability to regulate photosynthesis was assessed by setting up a protocol where dark-adapted leaves were exposed to an increasing light intensity up to 2000 μmol photons m^-2^ s^-1^. Light was then switched off monitoring relaxation in the dark for further 20 minutes (Figure 2). During this light treatment, several parameters describing the properties of both PSI and PSII and their ability to activate photo-protection responses like NPQ were measured using a DUAL-PAM-100 (Table 1). Plant response to increasing illumination intensity is a complex phenomenon and in order to describe it quantitatively NPQ induction curves were fitted with a sigmoidal function found to be the most suitable to describe the kinetics, similarly to what was previously observed in algae (Bernardi *et al*., 2016) (Figure S3).

**Figure 2.**
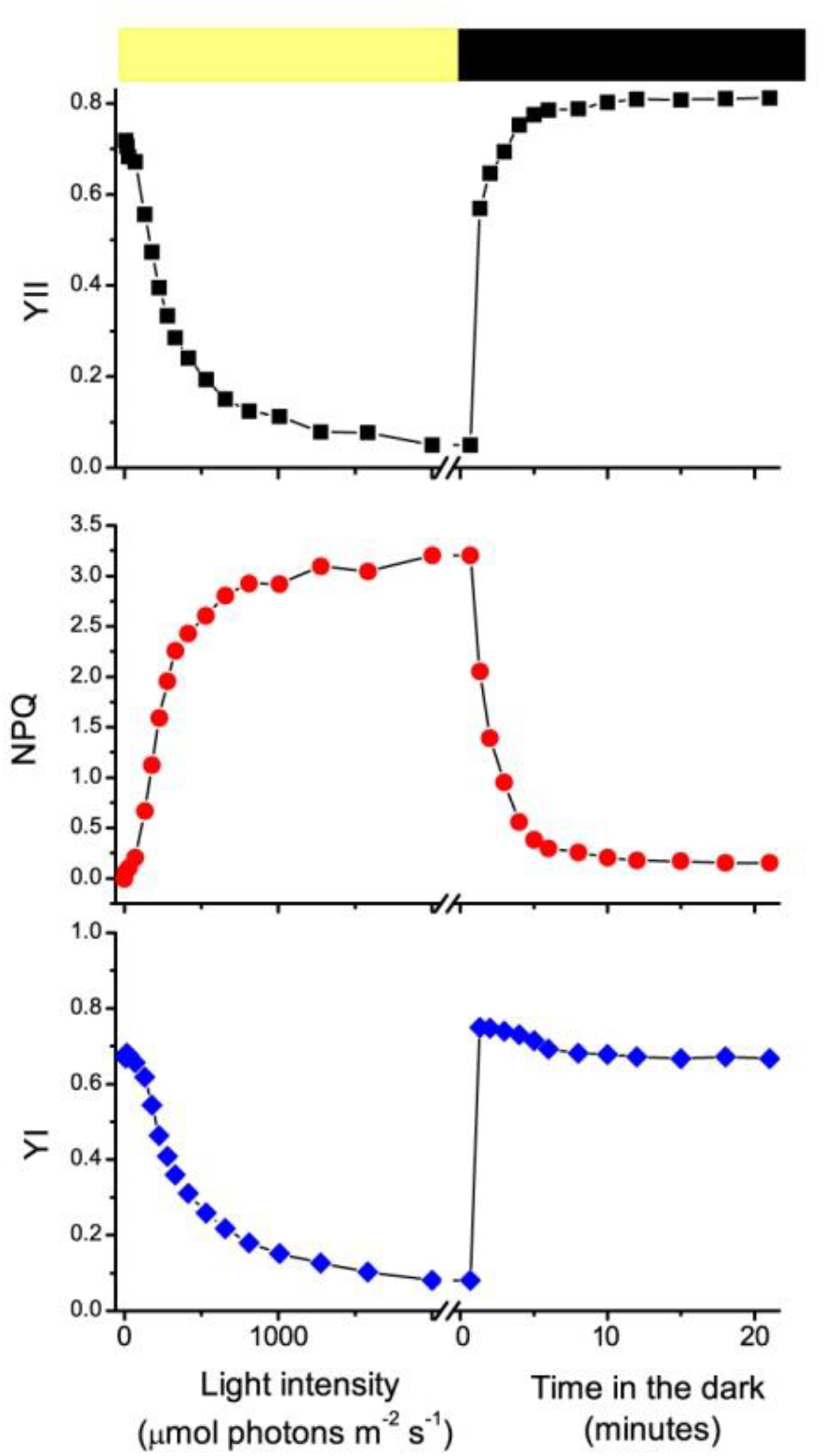
Representative measurements of plant photosynthetic performances. Dark adapted leaves were exposed to increasing light intensities (60 seconds for each point, 17 total minutes), followed by 20 minutes of dark recovery. Examples of measurements of YII (black squares), NPQ (red circles) and YI (blue diamonds) are shown from a *Taxus baccata* leaf measured in November 2016. Light and dark exposition are indicated with a yellow and a black bar.

Satisfactory fittings were obtained for all samples, yielding the quantification of three parameters NPQ_max_, k_NPQ_ and n_NPQ_, where NPQ_max_ is the maximal value reached, k_NPQ_ is the light intensity needed to reach 2/3 NPQ_max_ and n_NPQ_ is a parameter describing the sigmoidicity of the curve. The same approach was also used for PSI efficiency (YI), again obtaining a good description of the experimental curve recorded (Table 1). A bi-exponential decay was instead employed to obtain a quantitative description of NPQ relaxation in the dark, following previous examples (Nilkens *et al*., 2010; Holzwarth *et al*., 2013). These analyses yielded 5 parameters describing the kinetics: the amplitude and time constants of the two exponential components and a residual value (A1_qE_, A2_qZ_, t1_qE_, t2_qZ_ and y0_qI_ respectively) as reported in table 1 (Figure S3).

**Table 1.** List of parameters considered for the analysis. Each measurement yielded several parameters that were collected for each sample, describing environmental conditions during sampling, and PSII properties.

|  | Parameter | Description |
| --- | --- | --- |

| Environmental | Temp | Average temperature |
| --- | --- | --- |
|  | Light | Maximum light recorded |
| PSII | $F_v/F_m$; $ETR_{II_{max}}$; $Y_{II_{min}}$; $Y_{II_{recovery}}$; | PSII efficiency in the dark; Maximal $ETR_{II}$ value; minimal value for PSII efficiency under illumination; $Y_{II}$ after 20 minutes dark recovery |
| | $k_{NPQ}$, $n_{NPQ}$, $NPQ_{max}$ | parameters obtained fitting NPQ induction with a sigmoidal curve (see Figure S3) |
| | $A1_{qE}$, $A2_{qZ}$, $t1_{qE}$, $t2_{qZ}$ and $y0_{qI}$ | parameters obtained fitting NPQ relaxation with an exponential curve (see figure S3) |
| PSI | $YI_{max}$; $ETRI_{max}$; $YI_{min}$; $YI_{recovery}$; $Y_{ND_{max}}$; $Y_{NA_{min}}$ | maximal $YI$ measured; Maximal value for $ETRI$; minimal $YI$ under strong illumination; $YI$ after 20 minutes dark recovery; maximal value for $Y_{ND}$; minimal value for $Y_{NA}$ |
| | $k_{YI}$, $n_{YI}$, $YI_{start}$, $YI_{end}$ | parameters obtained by fitting $YI$ dynamics under illumination with a sigmoidal curve |

The measurements described above were repeated several times in a 2-year timespan and considering that some species lost their leaves during wintertime, a total of approx. 150 measurements was accumulated. To analyse this dataset, the presence of statistically significant correlations between all different parameters was first assessed using a Pearson product-moment with a Bonferroni correction (Table S1). It is important to underline that this analysis is here used as a screening tool to analyse a large dataset and identify the presence of a significant correlation between two parameters, but this does not imply the existence of a mechanistic connection.

### Effect of environmental conditions on Photosystem II photoinhibition

The statistical analysis reported in Table S1 showed significant correlation between environmental and photosynthetic parameters. Temperature in fact showed a statistically significant correlation with F_v_/F_m_ of dark adapted leaves (p > 0.95). A punctual analysis of all experimental data in Figure 3A showed that in the 18-30°C range all plants were characterized by high PSII efficiency, while at lower temperatures several plants showed major reduction in F_v_/F_m_ even if some still maintained high PSII activity. Data in Figure 3A thus suggest that low temperatures facilitate photoinhibition but do not necessarily cause it.

**Figure 3.**
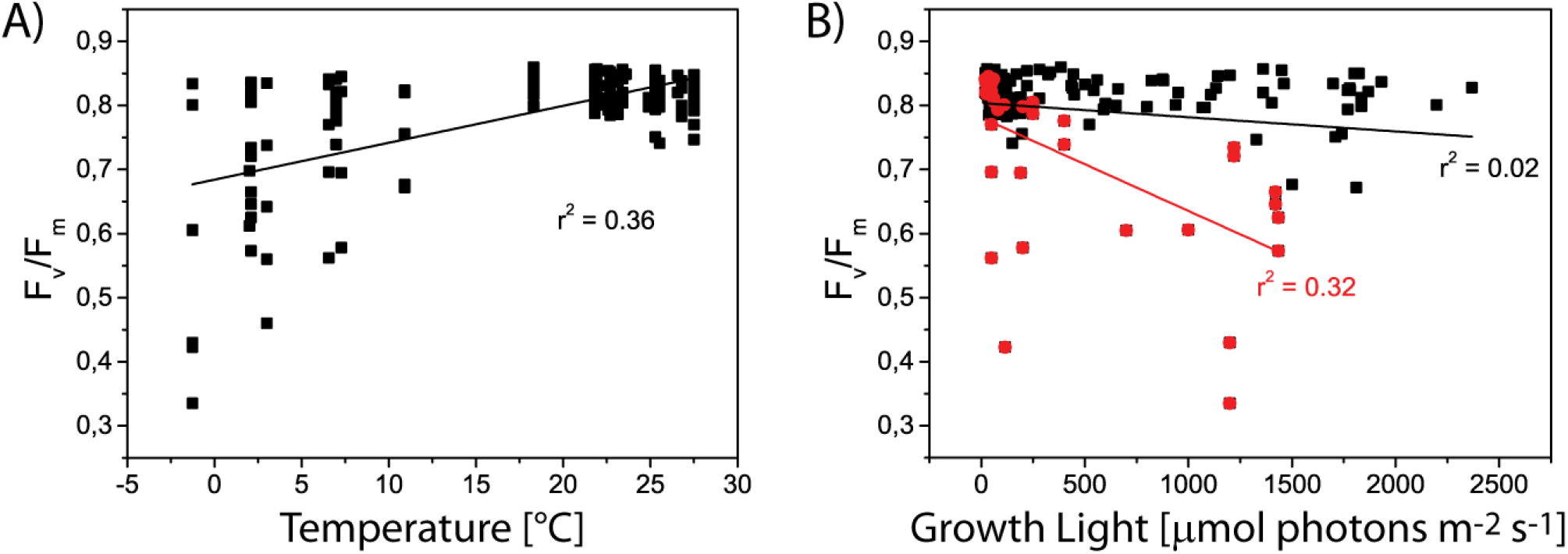
Effect of environment on maximum PSII efficiency. Relationship between temperature (A), growth light (B) and F_v_/F_m_. In (B), low temperature samples (< 10°C) are shown in red.

Another major parameter expected to influence photoinhibition is light intensity. Considering all data together there was no correlation between growing light and F_v_/F_m_, while a negative correlation emerged when only low temperature samples were considered (Figure 3B). These results thus confirm a well-known synergy between low temperatures and high light intensities in causing photoinhibition (HUNER *et al*., 1993), but also support the validity of our data analysis to identify potential correlations.

### Concerted acclimation of photosynthetic electron transport capacity to different light intensities

Light intensity is a major parameter for photosynthetic organisms and therefore it was repeatedly measured during sampling days. It should however be considered that light intensity can rapidly change and thus its measurement represents a rough estimation of average illumination experienced by the plants over multiple days. Despite this potential limitation light intensity monitored during sampling days showed several significant correlations with functional parameters, such as PSI and PSII maximal electron transport capacity (Figure 4A-B). This correlation suggests that all plants tend to respond similarly to a sustained stronger illumination by increasing their capacity of photosynthetic electron transport for both photosystems to ensure a higher ability to use available energy for photochemistry.

**Figure 4.**
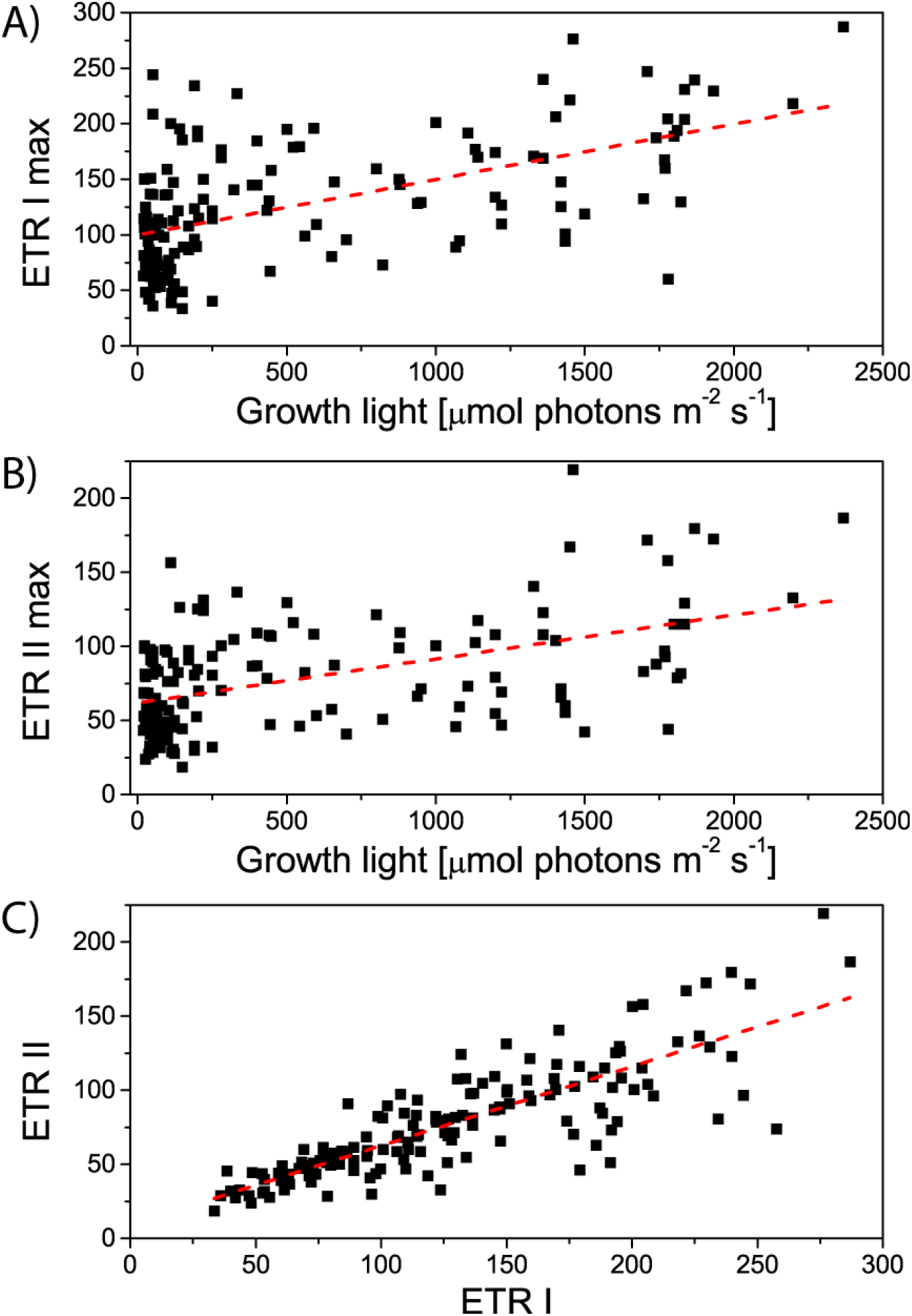
Correlation of growing light and maximum ETRI (A) and II (B). C) correlation between maximum ETRI and ETRII considering all point (black and red).

ETR I and II maximal values also show a strong correlation with each other (Figure 4C). This suggests that in different growing conditions all plants modulate concurrently PSI and PSII electron transport capacity, suggesting a strong tendency to balance the activity of the two photosystems.

### Variability and acclimation of NPQ

One of the major aims of the work was to sample NPQ variability between different plants (Jung & Niyogi, 2009; Oakley *et al*., 2018) also including effects due to acclimation to seasonal conditions. NPQ relaxation kinetics were fitted using a bi-exponential decay (Figure S3) to quantify three kinetic components, the fastest corresponding to qE, a slower one corresponding to qZ and the residual qI after 20 minutes of dark relaxation. Analysis of data obtained showed that time kinetics of different NPQ components (t_qE_ and tq_Z_) are not correlated with any other parameter and they do not respond significantly to different growth conditions.

Conversely, the analysis of the amplitude of the different components provide significant insights. qE, quantified from A_qE_, is the largest NPQ component and shows a strong correlation with NPQ max values (Figure 5A), differently from qZ and qI (Figure S4). The analysis of distribution of all components also shows clearly that A_qE_ is the major NPQ component in most cases, followed by A_qZ_ and A_qI_ (Figure 5B).

**Figure 5.**
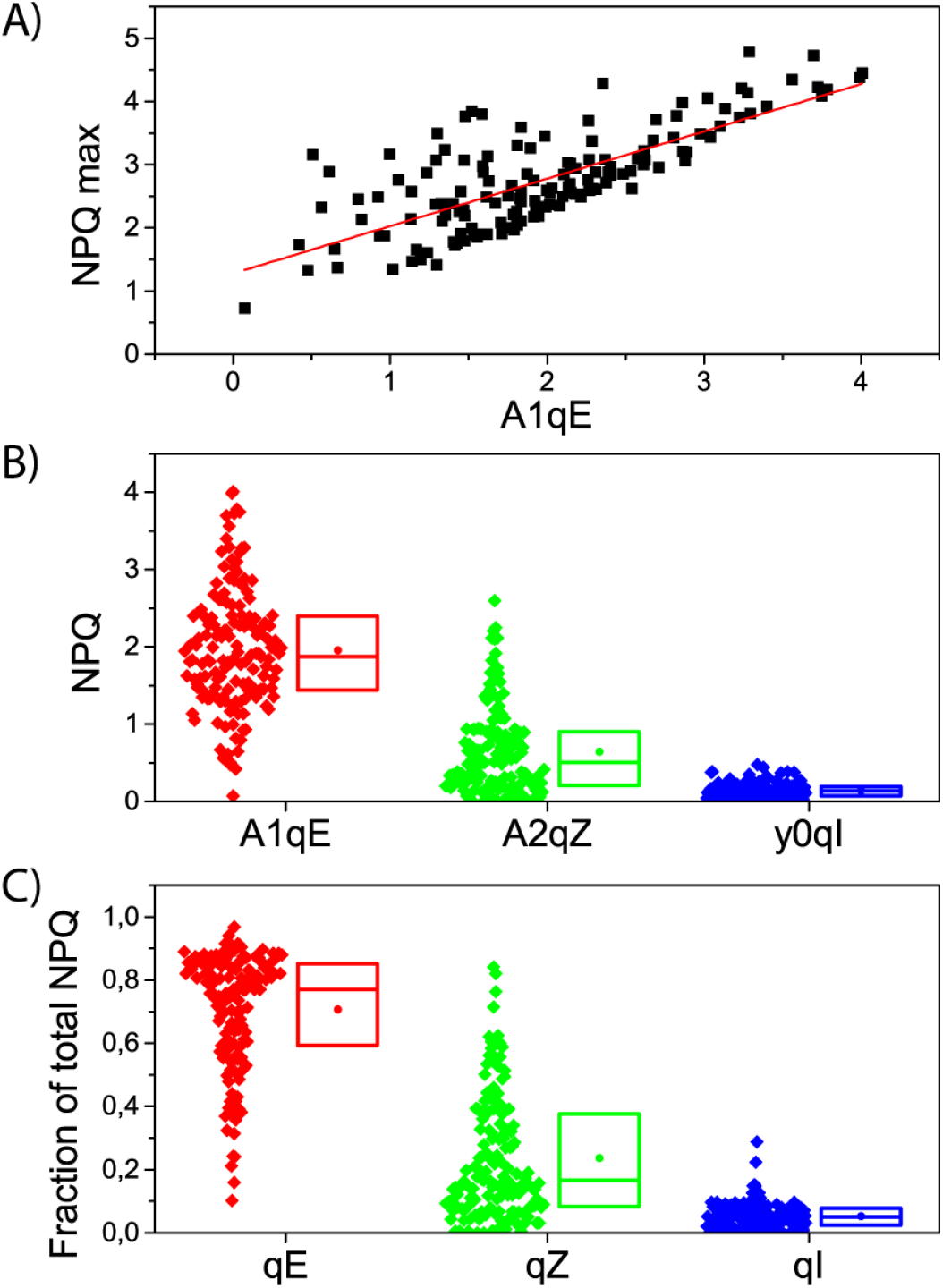
Contribution of different components to NPQ. A) Correlation between AqE and NPQ max. Correlations with AqZ and AqI are shown in Figure S4. B) distribution of relative amplitude of different NPQ components. Data are shown as diamonds on the left. On the right a box shows 25, 50 and 75 percentile distributions together with the mean (circle).

If the relative contribution of each component to total NPQ is considered (Figure 5C) this confirms that qE commonly represents at least 80 % of the total NPQ. At the opposite extreme qI contribution is always small and contributes to 0-10% of total NPQ (Figure 5C). qZ is commonly around 10-20% but in a few cases it can be much higher, reaching values as high as 80%, where, consequently qE contribution is much smaller. A qZ component is present in all plants measured but its absolute and relative contribution to NPQ is thus found the be the most variable.

Figure 6 shows examples of NPQ relaxation kinetics in plants where contribution from qE and qZ was extremely different, *Selaginella kraussiana, Selaginella pallescens, Taxus baccata* and *Gingko biloba*. As shown in Figure 6 the difference in qZ contribution affect not only NPQ intensity but also its kinetics. 30-500 seconds after the light is switched off (Figure 6A), in fact, plants with higher qZ have stronger residual NPQ than the others, suggesting that modulation of qZ is a possible strategy to regulate NPQ intensity but also its kinetics properties.

It is interesting to assess if this variable qZ is a species-specific property or if environmental conditions play instead a major role. To this aim the relative qZ contributions to total NPQ was analysed grouping values depending on the species (Figure 6B). These data show that acclimation has a significant influence on the relative amplitude of qZ and in some cases, e.g. species #8 or 9 *(Taxus baccata* and *Cephalotaxus harringtonii* respectively) there is a large difference in measurements performed in different times of the year. On the other hand, not all species behaved equally and, in some cases, qZ was never found to be a major component. The relative qZ contribution also does not correlate with temperature nor with growth light intensity (Figure S5). Overall these results suggest that plants can modulate qZ amplitude depending from growing conditions but likely this response is also influenced by species-specific features and it is not equivalent in all organisms.

**Figure 6.**
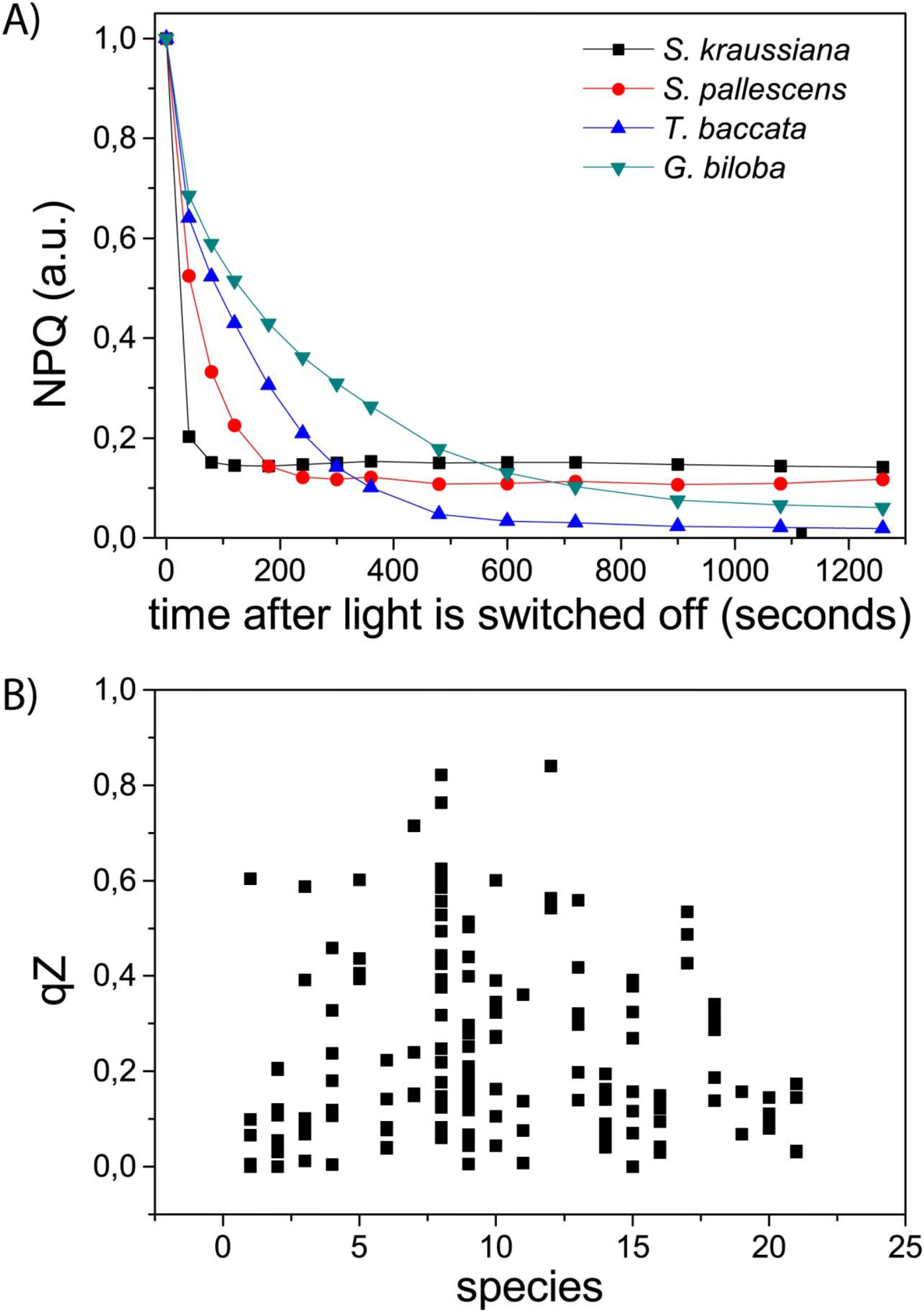
Variability in qZ contribution. A) Examples of NPQ relaxation kinetics with contrasting qZ component. NPQ was normalized to the maximal value before the light was switched off. Original data are in Figure S5. ; B) qZ contribution in different species, species number indicates the order from figure 1.

When measurements from the same species in different seasons are compared, it appears that in some cases NPQ intensity is modulated upon acclimation (see below). When all measurements are considered globally however, there is no correlation between the maximal NPQ value reached by each sample and growth light (Figure 7). On the contrary the two parameters seem completely independent, suggesting that there is no general tendency of species growing in more intense light to have a stronger NPQ.

On the contrary there is a small but statistically significant correlation between growth light and K_NPQ_, the light intensity needed to reach 50% of NPQ maximal value. This suggested that, overall, acclimation to different light intensities influence the light intensity values where NPQ is activated and in general if plants experience strong illumination they will acclimate and use light more efficiently and therefore activate NPQ at higher intensities.

**Figure 7.**
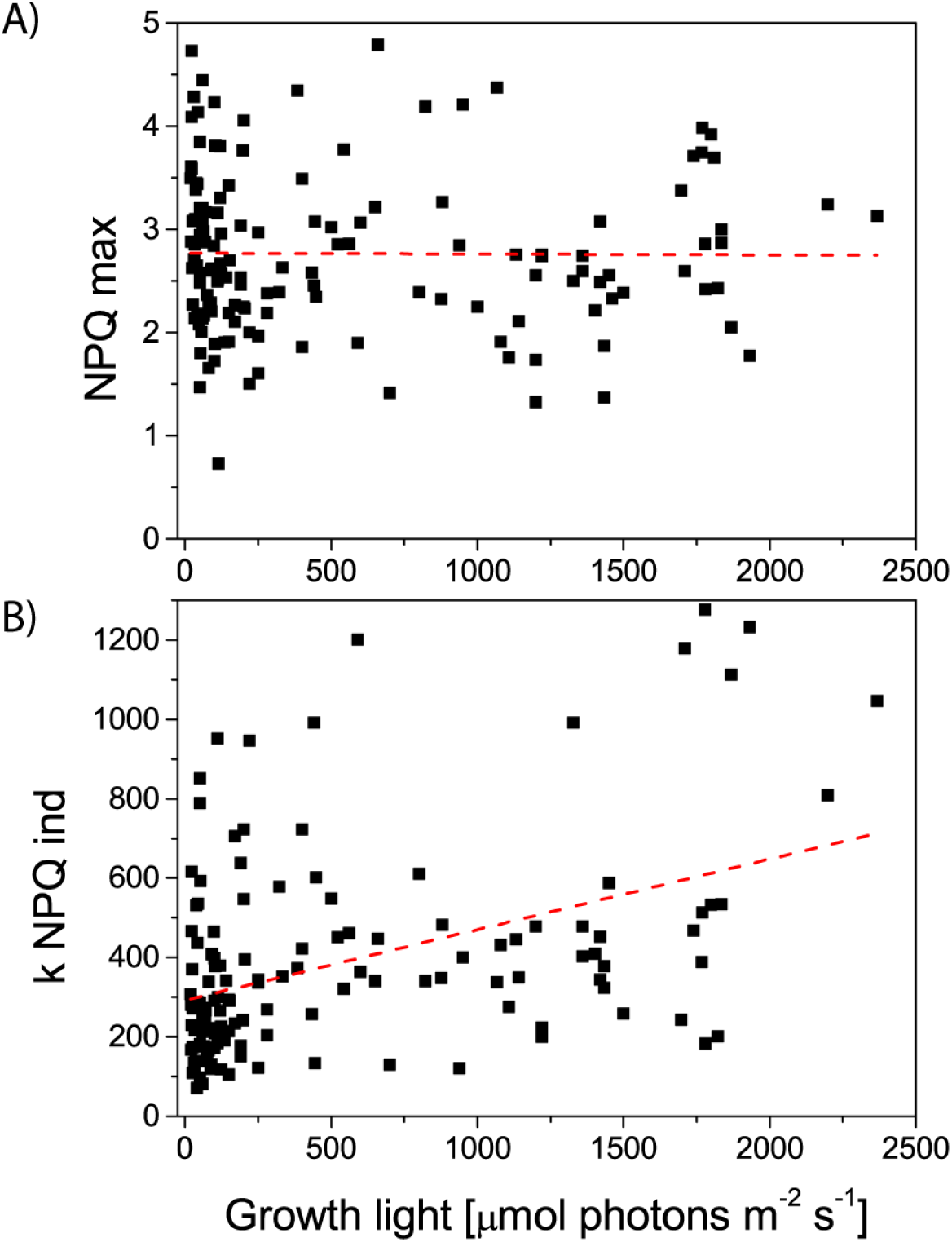
NPQ acclimation to different light intensities. A) correlation between NPQ max and growth light. B) Correlation of kNPQ with growth light, where NPQmax is the maximum NPQ value reached during the actinic phase while kNPQ is the intensity where NPQ reaches 2/3 of the maximal value.

Photoinhibition has also been suggested to participate to protection by contributing to heat dissipation quantifiable from NPQ with its slower component, qI, that was correlated with the accumulation of inhibited PSII. qI was here estimated from the Y_0_ NPQ parameter that quantifies NPQ with a very slow relaxation. Y_0_ was found variable, going from 0 to 0.47, as shown in example of kinetics reported in Figure S5. qI value, however, did not correlate with the level of photoinhibition suffered during growth nor with factors influencing photoinhibition like light intensity (Figure 8). This suggests that ability to activate qI is not correlated with photoinhibition experienced during growth but is rather a specific response activated by illumination during NPQ measurement.

**Figure 8.**
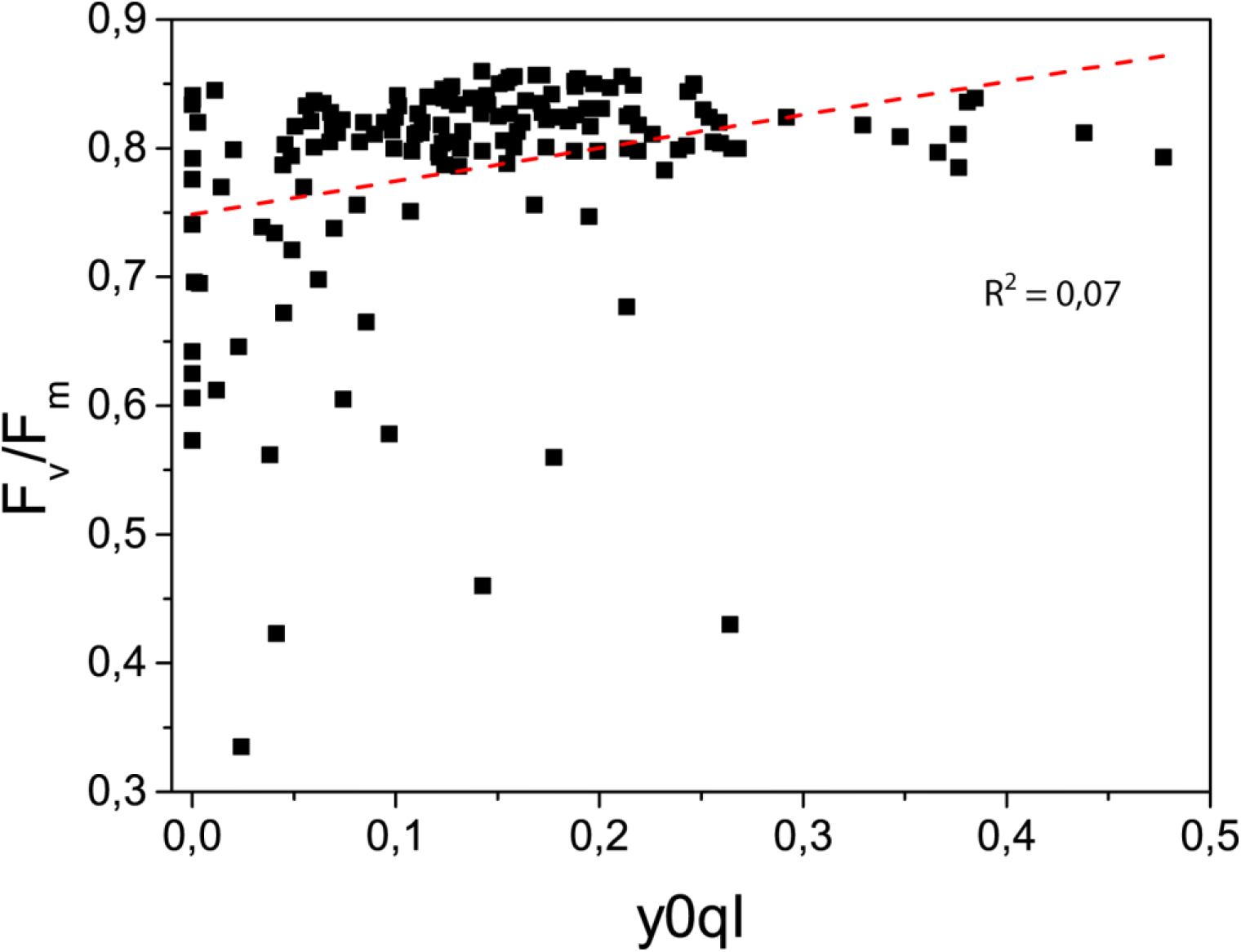
Effect of environmental parameters on photosynthetic properties. Correlation between qI and Fv/Fm.

### Seasonal acclimation of an evergreen plant

In the pool of selected species there were some evergreen plants that faced seasonal acclimation to low temperatures and/or high solar radiation. As example, *Taxus baccata* NPQ activation kinetics for the former are shown in figure 9. Light exposed leaves clearly show a modulation of NPQ activation, with some leaves requiring more intense illumination. NPQ max is also modulated (Figure 9C) during the seasons with wintertime leaves showing a tendency to have higher values.

**Figure 9.**
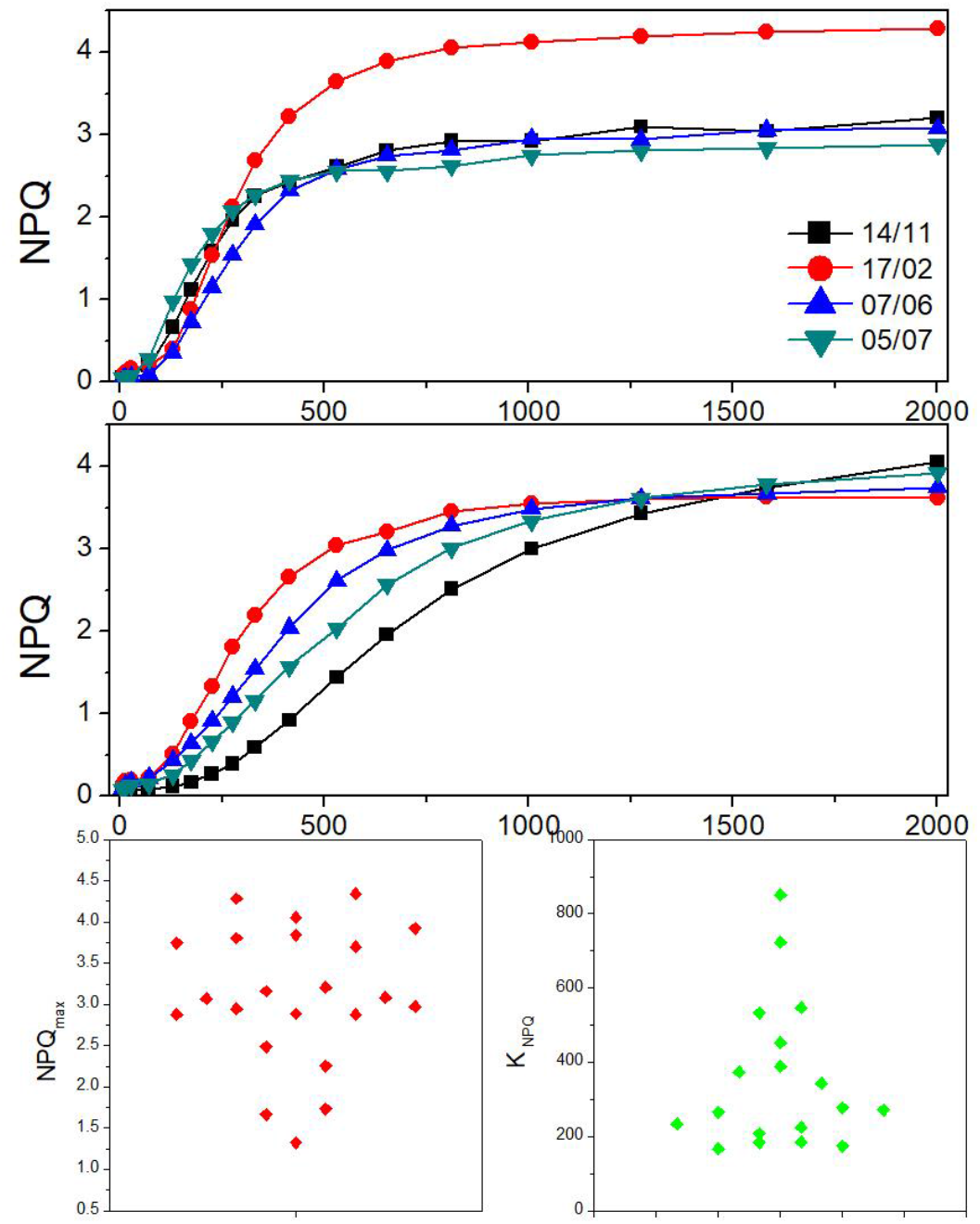
NPQ Acclimation of evergreen plant. *Taxus baccata* leaves were measured multiple times in different seasons. Example of NPQ induction with an increasing light treatments are shown. A) examples of shade adapted leaves. B) example light exposed leaves measured. As example measurements performed the same four days (one for each season) are shown. C) NPQmax values for all measurements for this species; D) K NPQ values.

## DISCUSSION

The present work explored the ability of 20 different species to activate photosynthesis regulatory responses, sampling in different seasons and canopy position to assess acclimation influence. The measurements yielded several parameters describing photosynthetic properties of the plants that were analysed (table 1 and S1) looking for common trends and peculiarities. These analyses yielded some expected results, such as that low temperature in combination with high illumination increases the probability of photoinhibition, supporting the validity of the approach.

### Acclimation vs. adaptation

One objective of the work was to assess regulation of photosynthetic apparatus responses showed correlation with plants evolutionary position (ferns, gymnosperms, angiosperms). In the dataset analysed there was no specific correlation emerging between plants phylogenetic groups and any photosynthetic response. This suggests that regulatory mechanisms involved in the responses assessed were all present in common ancestors of species analysed. This is well consistent with the available information on the molecular mechanism responsible of mechanisms like NPQ or xanthophyll cycle that are known to be conserved in all plants (Li *et al*., 2000; Gerotto & Morosinotto, 2013).

Acclimation to different growing conditions such as light intensity and temperature were found instead to have a larger influence on the photosynthetic properties. Some common trends were observed, the most evident being the correlation of growing light with both ETR I and ETR II. In all plants analysed here not only PSI and PSII activity respond similarly to growing light but their electron transport ability was found to have a strong correlation (Figure 4C). This suggest the presence of a strong advantage in avoiding unbalances in electron transport activity of the two photosystems.

### Modulation of Non Photochemical quenching activity and kinetics is a species specific trait

Non photochemical quenching is a fast response to changes in illumination that is activated in seconds and is important for plants growth under light fluctuation (Külheim *et al*., 2002). Despite being a short-term response NPQ was also shown to be modulated upon acclimation in different growing light conditions. As example, in *Arabidopsis thaliana*, the moss *Physcomitrella patens* or even in the green alga *Chlamydomonas reinhardtii* the intensity of NPQ correlated with light intensity during growth (Ballottari *et al*., 2007; Peers *et al*., 2009; Gerotto *et al*., 2011). In this work as well, we observed cases (see Figure 9) where NPQ was increased upon acclimation to stronger illumination. However, there was no general trend of plants exposed to stronger illumination to have a higher NPQ (Figure 7A). This is not due to a lack of sampling plants in different acclimation states since responses in ETR modulation are clearly visible and modulation of K_NPQ_ is also detected (Figure 7B). On the contrary this is a clear indication that different species can respond differently in their NPQ modulation (Gerotto & Morosinotto, 2013). Photosynthetic organisms possess a large set of regulatory mechanisms to respond to different environmental conditions (Eberhard *et al*., 2008) and the relative weight that NPQ plays in this portfolio of responses is variable and depends from the specific species and environmental niche.

The exploitation of the variability of NPQ responses also showed that plants also have the possibility of modulating the kinetic properties. A possible strategy to achieve this objective is the modulation of relative contribution of qZ component. Different species in fact show a variable contribution of this component and, consequently, they have different NPQ kinetics, as exemplified in figure 6. This ability to modulate NPQ kinetics could be important to respond to different conditions where not only the light intensity is variable but also the frequency of dynamic changes can change. Indeed the kinetic properties of NPQ have been shown to be particularly important for crops photosynthetic productivity (Kromdijk *et al*., 2016).

Finally, another major information is that contribution to qI component to total NPQ is found constant in all species and growing conditions suggesting a limited regulation of this phenomenon.

It is also interesting that this does not correlate with photoinhibition suffered by plants during growth, suggesting that despite the name other mechanisms (e.g. (Malnoë *et al*., 2017)) may play a major role in this slow NPQ component.

